# Genetic inactivation of SARM1 axon degeneration pathway improves outcome trajectory after experimental traumatic brain injury based on pathological, radiological, and functional measures

**DOI:** 10.1101/2021.03.07.434264

**Authors:** Donald V. Bradshaw, Andrew K. Knutsen, Alexandru Korotcov, Genevieve M. Sullivan, Kryslaine L. Radomski, Bernard J. Dardzinski, Xiaomei Zi, Dennis P. McDaniel, Regina C. Armstrong

## Abstract

Traumatic brain injury (TBI) causes chronic symptoms and increased risk of neurodegeneration. Axons in white matter tracts, such as the corpus callosum (CC), are critical components of neural circuits and particularly vulnerable to TBI. Treatments are needed to protect axons from traumatic injury and mitigate post-traumatic neurodegeneration. The *Sarm1* gene is a central driver of axon degeneration through a conserved molecular pathway. *Sarm1*-/- mice with knockout (KO) of the *Sarm1* gene enable genetic proof-of-concept testing of *Sarm1* inactivation as a therapeutic target. We evaluated *Sarm1* deletion effects after TBI using a concussive model that causes traumatic axonal injury and progresses to CC atrophy at 10 weeks, indicating post-traumatic neurodegeneration. *Sarm1* wild-type (WT) mice developed significant CC atrophy that was reduced in *Sarm1* KO mice. Using electron microscopy to quantify individual axons demonstrated that *Sarm1* KO preserved more intact axons and reduced damaged or demyelinated axons. MRI in live mice identified significantly reduced CC volume after TBI in *Sarm1* WT mice that was attenuated in *Sarm1* KO mice. MR diffusion tensor imaging detected reduced fractional anisotropy in both genotypes while axial diffusivity remained higher in *Sarm1* KO mice. Immunohistochemistry revealed significant attenuation of CC atrophy, myelin loss, and neuroinflammation in *Sarm1* KO mice after TBI. Functionally, TBI resulted in late-stage motor learning and sleep deficits that were ameliorated in *Sarm1* KO mice. Based on these findings, *Sarm1* inactivation can protect axons and white matter tracts to improve translational outcomes associated with CC atrophy and post-traumatic neurodegeneration.

## INTRODUCTION

Traumatic brain injury (TBI) results in long term disability in more severe cases and can cause persistent symptoms even in patients who receive a “mild” diagnosis [41, 64, 69]. TBI may also lead to post-traumatic neurodegeneration and increase the risk for co-morbid neurodegenerative diseases, such as Alzheimer’s disease [15, 18, 30, 70]. In patients with moderate-severe TBI, diffuse axonal injury has been shown to predict the extent of post-traumatic neurodegeneration, based on MRI volumetric and diffusion tensor imaging (DTI) data [29]. The strongest relationship was found in central WM tracts, including the corpus callosum (CC). The CC is one of the main structures exhibiting atrophy across patients with complicated mild to severe TBI [15, 30]. Furthermore, DTI tractography demonstrated disrupted fiber tract continuity in anterior CC regions after concussions while broad areas of disrupted tracts were found throughout the CC in patients with MRI findings of diffuse axonal injury [37]. *Ex vivo* MRI of a patient who died 26 days after TBI demonstrated that white matter tract disruptions detected by DTI correlated with neuropathological identification of axonal injury [66].

Long axons in white matter tracts are particularly vulnerable to damage in all forms of closed head TBI [10, 39, 55]. At an early stage of damage, axons can recover; once mechanical or molecular processes fragment the axon, then the distal axon irreversibly degrades by Wallerian degeneration [31, 36, 53, 54, 65, 89]. The majority of axon damage is due to secondary mechanisms of damage that can cause axons to initiate Wallerian degeneration for weeks or more after the initial TBI event [12, 53, 54]. Irreversible axonal injury leads to disconnection between brain regions and disruption of neural circuits that may contribute to diverse TBI symptoms [32]. To improve outcomes for patients after TBI, research is needed to identify approaches to protect against axon degeneration and, further, to determine whether acute axon protection can reduce post-traumatic neurodegeneration.

Wallerian degeneration is an “active program of axon self-destruction” [73]. After injury or mitochondrial dysfunction, the SARM1 (sterile alpha and Toll/interleukin-1 receptor motif-containing 1) protein executes a highly conserved molecular axon death pathway [26, 47, 67, 80]. Injury releases SARM1 from an auto-inhibited state that is found in healthy axons [21, 73, 76]. Active SARM1 is a glycohydrolase which depletes nicotinamide adenine dinucleotide (NAD+) that is critical for energy stores in axons [17, 22, 25]. Loss of sufficient NAD+ leads to a cascade of axon destruction in which ionic imbalance leads to cytoskeletal and structural breakdown within the axon [24, 25, 81].

Deletion of the *Sarm1* gene provides a genetic “proof-of-concept” to examine the effects of long-term inactivation of this axon death pathway in chronic TBI and to evaluate the relationship of axonal injury to post-traumatic neurodegeneration. Knockout of *Sarm1* in cultured human sensory neurons and in live mice reduced axon damage in trauma and in chemotherapy-induced models of peripheral neuropathy [13, 23, 25, 67]. In multiple models of TBI, *Sarm1* knockout (KO) mice exhibited significantly reduced axon damage at acute and late phase time points [34, 51, 56, 92] Long-term studies of TBI in mice can model post-traumatic neurodegeneration that includes atrophy of the CC [46, 52, 61]. Initial results from our prior studies provided the first evidence that *Sarm1* deletion may reduce both late stage axon damage and CC atrophy [51].

We now focus on CC atrophy as an important outcome measure of post-traumatic neurodegeneration. We evaluate the effect of *Sarm1* deletion on axon damage, demyelination, and neuroinflammation on the progression of CC atrophy after closed head TBI. We use longitudinal MRI to detect changes in CC volume and white matter integrity in live mice, which provides a highly translational outcome measure of prolonged *Sarm1* inactivation. To demonstrate whether the pathological effects of *Sarm1* deletion have a meaningful impact on complex functions, we assess motor skill learning and sleep behavior. This experimental design provides a rigorous and translationally relevant screen of *Sarm1* genetic inactivation as a potential therapeutic strategy for improving outcome measures after TBI.

## MATERIALS AND METHODS

### Mice

All mice were treated in accordance with guidelines of the Uniformed Services University of the Health Sciences and the National Institutes of Health Guide for the Care and Use of Laboratory Animals. Mice were socially housed with 2–5 mice per 35 cm × 16.5 cm × 18 cm cages containing enrichment objects. Mice were maintained on a standard 12 h cycle of daytime light (6:00–18:00). *Sarm1-/-* knockout (KO) mice, B6.129 × 1-Sarm1tm1Aidi/J (RRID:IMSR_JAX:018069) were originally obtained from The Jackson Laboratory. *Sarm1* KO mice were crossed to C57BL/6J mice and then heterozygotes were bred to generate littermates for experiments. *Sarm1* is highly expressed in the brain; lack of detectable SARM1 protein in *Sarm1* KO mice does not produce gross or microscopic brain pathology [27, 42]. The colony was maintained at Charles River Laboratories (Wilmington, MA) where ear biopsies were genotyped using a 3-primer allele specific PCR assay that targets the mutated *Sarm1* region. Experimental mice were acclimated for 3 days after shipment and prior to the start of experiments. The total number of mice used in experiments was *Sarm1* WT (n = 66; 33 male, 33 female) and *Sarm1* KO (n = 75; 34 male, 41 female) littermates.

### TBI and Sham Procedures

The TBI model has been characterized in our previous studies [51, 58]. This concussive, closed head injury model results in pathology under the impact site at bregma in the CC and the adjacent cingulum [52, 79]. Under 2% isoflurane anesthesia, 8–10 week old mice received a single impact onto the skull at bregma using an ImpactOne stereotaxic impactor (Leica Biosystems, Buffalo Grove, IL) with a 3-mm-diameter tip (velocity set at 4.0 m/s; depth of 1.5 mm; dwell time of 100 ms). Sham mice received the same procedure without the impact. Righting reflex demonstrated a significant injury effect of longer time to righting after TBI, as compared to sham procedures, which did not differ with *Sarm1* genotype or sex (Figure S1, Online Resource 1). The predetermined study design criteria required exclusion of three mice for depressed skull fracture and/or impactor malfunction.

### Electron Microscopy

Transmission electron microscopy (EM) was used to analyze axon and myelin subcellular structure, which can reveal a broad range of axon and myelin pathology [52, 58]. Mice were anesthetized with ketamine/xylazine before transventricular cardiac perfusion with 4% paraformaldehyde (Electron Microscopy Sciences, Hatfield, PA; Cat #19210) and 2.5% glutaraldehyde (Electron Microscopy Sciences; Cat #16210) in 0.1 M phosphate buffer. After overnight post-fixation, brains were cut into sagittal (40 μm) sections using a Leica VT-1200 vibrating blade microtome (Leica Biosystems, Buffalo Grove, IL). Parasagittal sections were immersed in 2% osmium tetroxide (OsO4; Electron Microscopy Sciences; Cat #19100) infiltrated with Spurr epoxy resin (Electron Microscopy Sciences; Cat #14300), flat-embedded and then polymerized at 70 °C for 11 h. Thin sections (∼70 nm) were cut on an Ultracut UC6 ultramicrotome (Leica Biosystems). Copper grids containing thin sections were post-stained for 20 min in 2% aqueous uranyl acetate (Electron Microscopy Sciences; Cat #22400) and for 5 min in Reynolds lead citrate (Reynolds, 1963).

### Quantification of sagittal CC width and electron microscopy analysis

Cohorts consisted of Sarm1 WT sham (n = 9; 4 female, 5 male) and TBI (n = 8; 4 female, 4 male) along with *Sarm1* KO sham (n = 7; 3 female, 4 male) and TBI (n = 9; 5 female, 4 male). Mice were sacrificed at 10 weeks after sham or TBI procedure. Resin-embedded 40 μm sagittal sections approximately 200 μm lateral to the midline were osmicated to stain myelin and imaged in bright field on an Olympus IX-70 microscope. CC width was measured across the superior to inferior borders in five locations at ∼100 μm intervals across a 0.5 mm rostro-caudal region centered under bregma. This region-of-interest (ROI) consistently exhibits traumatic axonal injury in this TBI model, as evidenced by dispersed damaged axons among adjacent intact axons, and matches the EM analysis performed in our previous work with this *Sarm1* line of mice [51, 52].

Thin sections for EM analysis were then cut from within the CC ROI of the 40 μm thick sections. The EM grids of sagittal thin sections were reviewed on a JEOL JEM-1011 transmission electron microscope (JEOL USA Inc., Peabody, MA) and images were acquired using an AMT XR50S-A digital camera (Advanced Microscopy Techniques, Woburn, MA). Images were taken at 5000× magnification and 8-10 images per animal were quantified for classification of axon and myelin pathology. For each image, a 17 μm × 12.5 μm region defined the counting frame, within which >120 axons were quantified. All axons within the counting frame were counted. Axons partially crossing the top and right lines were included while those partially crossing the left and bottom line were excluded. Axons were classified as intact axons, de/unmyelinated axons, axons with abnormal mitochondria, or damaged axons [51, 52, 58]. Damaged axons were defined as axons with cytoskeletal disruption or axons with accumulated vesicles and debris. Mitochondria that appeared swollen encompassed > 50% of the area of the axon cross-section and were considered abnormal. De/unmyelinated axons were > 0.3 µm in diameter and lacked detectable compact myelin, but otherwise appeared intact. Axons without myelin and with a diameter < 0.3 µm were excluded as this axon size is typically unmyelinated in the CC of healthy adult mice [77]. TBI-induced demyelination was inferred when de/unmyelinated axon counts were significantly greater after TBI, as compared to sham. Myelin outfoldings were identified as myelin extending out from an axon and folding back onto itself to form double layered or redundant myelin [58] but were not significantly induced by TBI at this 10 week time point in mice of either genotype (data not shown). Additional images were taken at 10,000–15,000x for illustration of pathological features.

### Magnetic resonance imaging (MRI) analysis of CC volume and microstructure

MRI with multi-spin-echo, high resolution T2-weighted (T2-w) and diffusion tensor images (DTI) were acquired to assess the longitudinal effects of *Sarm1* deletion in the CC ROI following TBI. Longitudinal MRI studies were conducted in live mice with repeated scans of the same mouse at baseline (prior to TBI) and at 3 days and 10 weeks post-TBI. Mice were scanned on a 7 tesla Bruker BioSpec 70/20 USR Magnetic Resonance Imaging System with a 660 mT/m, 12 cm diameter gradient coil, 86 mm quadrature TX/RX coil, and Bruker Mouse head 4 channel receive coil array (Bruker BioSpin GmbH, Reinstetten, Germany). Mice were anesthetized in an induction chamber with a mixture of 4% isoflurane in medical air and maintained with 1.5–1.75% isoflurane in medical air delivered by nose cone during the MRI procedures. Respiration rate (range 40–70 BPM, maintained by adjusting isoflurane concentration) and temperature were continuously monitored throughout the experiments. MRI slices were positioned using a sagittal localizer so that coronal slices were oriented perpendicular to the length of the CC axis and each brain was aligned with the midline crossing of the anterior commissure within in the same coronal slice [78, 83, 91].

A whole brain T2 relaxation time map was generated from a two-dimensional rapid acquisition with relaxation enhancement (2D RARE, coronal [33] using the following parameters: TR = 4000 ms, echo time (TE) = 10, 30, 50, 70, 90, 110 ms, echo train length (ETL) = 2, number of averages NA = 4, field of view (FOV) 14 mm × 14 mm, matrix 112×112, in-plane resolution 125 µm × 125 µm, slice thickness 750 µm, number of slices NS = 21, no fat suppression, BW 36 kHz, time 14:56 min. Additionally, a high resolution proton density weighted (PD-w) 3D MRI was acquired to measure CC volumetrics using the following parameters: 3D RARE, TR = 2500 ms, TE = 30 ms, ETL = 8, number of averages NA = 1, FOV 18 mm × 14 mm × 11 mm, matrix 144×112×88, 125 µm isotropic resolution, no fat suppression, BW 36kHz, time 55:25 min. DTI data was acquired using the following parameters: 2D 2-shot echo planar imaging (EPI), TR = 3000 ms, TE = 27, NA = 2, 4 B0 and 30 non-collinear diffusion directions, b = 600, 1200 s/mm^2^, δ = 5 ms, Δ = 12 ms, FOV 14 mm × 14 mm, in-plane resolution 175 µm × 175 µm, matrix 80×80, slice thickness 750 µm, number of slices NS = 21, fat suppression, BW 300kHz, time 12:48 min.

T2-w images were converted to the NIFTI file format using a custom script in Matlab. A bias field correction was performed on the 30 ms echo time image using the N4BiasFieldCorrection command in the Advanced Normalization Toolkit (ANTs) [7]. The computed bias field was then applied to the other echo times (10, 50, 70, 90, 110 ms). Estimates of T2-decay (T2) and amplitude (S_0_) images (maps) were obtained using a nonlinear fit to the equation S_i_(TE) = S_0_* exp (−TEi/T2), where Si is the signal intensity for echo time TE_i_. The T2-map and amplitude values were fed into a deep learning algorithm to create a brain mask [71, 78, 91]. Brain masks were manually corrected as needed. The ROI in the CC was drawn manually using VivoQuant software (inviCRO, Boston, MA). The CC ROI was defined as extending from the midline bilaterally to the point of ventral curvature in the external capsule (Online Resource 2), as in our previous studies [78, 91].

DTI were processed using the TORTOISE v3.2.0 software package [62, 68]. Motion correction, eddy current correction, and EPI distortion correction were performed using the DIFFPREP function. Tensors were computed using a nonlinear tensor fit with RESTORE, and fractional anisotropy (FA), trace (TR), axial diffusivity (AD), and radial diffusivity (RD) were computed from the diffusion tensor images. An ROI was drawn manually in the CC within the slice under the impact site at bregma (Figure S2, Online Resource 1), and average CC values of FA, TR, AD, and RD were computed. TR did not show an effect of injury or genotype (data not shown).

To calculate volume change in the CC the 125 µm isotropic 3D T2-weighted image slices were converted to NIFTI file format using a custom script in Matlab. A bias field correction was performed using the *N4BiasFieldCorrection* command in the Advanced Normalization Toolkit (ANTs) [7]. The brain mask from the multi-spin-echo T2 image set was transformed to the 3D T2-w imaging using rigid registration and linear interpolation and thresholded at 0.5 to create a new binary mask. A template was created from the baseline scans using the antsMultivariateTemplateConstruction2.sh script [8]. Each image for each time point was then registered to the template using the nonlinear registration algorithm in ANTs [7]. Registration parameters were selected based on the approach described by Anderson et al. [3]. Voxel-wise maps of volume change were created using the *CreateJacobianDeterminateImage* function in the ANTs toolkit. A ROI was drawn manually in the CC on each of seven 125 µm coronal image slices (Online Resource 2) that encompassed the CC over the lateral ventricle and under the site of injury within the +0.5 and −0.5 mm window relative to bregma. The average value of volume change was then computed. The statistical parametric mapping (SPM) v12 software was used to perform a voxel-wise analysis of the volume change images.

### Immunohistochemistry

At 10 weeks following TBI or sham procedure, *Sarm1* mice were perfused with 4% paraformaldehyde and brains cut as 14 μm-thick coronal cryosections for immunohistochemistry. Myelin was detected by immunolabelling for myelin oligodendrocyte glycoprotein (MOG; polyclonal mouse anti-MOG; 1:100; Millipore, Burlington, MA; Cat# MAB5680, RRID: AB_1587278). Astrocytes were evaluated by immunostaining for glial fibrillary acidic protein (GFAP; monoclonal mouse anti-GFAP; 1:500; Millipore, Burlington, MA; Cat# MAB3402, RRID:AB_94844). Microglia/macrophages were identified using polyclonal rabbit antibody against ionized calcium binding adaptor molecule 1 (IBA1; 1:500; Wako, Richmond, VA; Cat# 019-19741, RRID:AB_839504). All tissue sections were counterstained with DAPI nuclear stain (Sigma-Aldrich, St. Louis, MO; Cat# D9542).

### Immunohistochemical analysis of coronal CC width, myelination, and neuroinflammation

Immunohistochemical analysis used cohorts of *Sarm1* littermates perfused at 10 weeks after the TBI or sham procedure for *Sarm1* WT sham (n = 7; 3 female, 4 male) and TBI (n = 7; 3 female, 4 male) along with *Sarm1* KO sham (n = 6; 2 female, 4 male) and TBI (n = 9; 5 female, 3 male). Images within the CC ROI were acquired with a 10x objective on an Olympus IX-70 microscope using a SPOT RT3 camera. For quantification in coronal images, the CC ROI extended from the midline laterally to under the peak of the cingulum at coronal levels between +0.5 and − 0.5 mm relative to bregma. The CC width (superior-inferior thickness) was measured as the average of measurements taken at the midline and bilaterally at ∼200 μm lateral to the midline, under the peak of the cingulum, and ∼200 μm lateral to the peak of the cingulum using MOG staining (Online Resource 2). ImageJ software was used to threshold fluorescence levels to quantify the area of immunolabeling above background [5, 51]. Images were also acquired with a 40x objective to classify the morphology of IBA1 immunolabeled cells as resting or activated cells [58, 79, 91]. Quantification included 4–6 sections per mouse.

### Motor skill learning task

The Miss-step wheel motor assay was performed using a protocol previously described [78]. The Miss-step wheel motor assay has been shown to engage the CC and be sensitive to changes in myelination [35, 57, 59]. From 8 to 10 weeks post TBI, mice were singly housed in home cages with a Miss-step running wheel equipped with an optical sensor to detect wheel revolutions (Mouse Miss-step Activity Wheel system, Cat#80821, Lafayette Instruments, Lafayette, IN). The Miss-step running wheels have 16 rungs missing from a standard wheel so that the remaining 22 rungs are distributed in an irregular interval pattern [35]. Mouse whiskers were clipped so that avoiding miss-steps was dependent on learning to follow the rung located on the prior step by bringing the hind paw forward to grasp the rung used by the forepaw [57]. Activity Wheel Monitor software (Lafayette Instruments) counted wheel revolutions at 6 min intervals during the “lights on” phase and 1 min intervals during the “lights off” phase. Results were exported to a Microsoft Excel file every 24 hrs.

### Sleep/wake pattern

Sleep pattern data was collected using a non-invasive automated scoring system (Signal Solutions LLC, Lexington, KY). Mice were single housed during the 8^th^ week following TBI or sham procedures for 72 hrs and maintained on a standard 12 h cycle of daytime light (6:00–18:00) with continuous data collection. A cage floor matt with piezoelectric sensors recorded 4 second epochs and used the 2-4 Hz breathing rhythm of mice to classify intervals of 30 seconds or more as asleep or awake [50]. This piezoelectric sensor system compares well with sleep data collected by visual observation and with electrophysiological discrimination of sleep/wake intervals, yet avoids the surgical procedures of electrophysiological techniques that could confound other assessments of TBI [60, 90].

### Statistical Analysis

Sample sizes were estimated for > 80% power based on prior data in *Sarm1* and in C57BL/6 mice [51, 91] (Figures S2, S3, Online Resource 1). Mice were randomized to TBI/sham using the RAND function in Microsoft Excel. Investigators involved in data collection and analysis were blinded to genotype and injury condition until after the conclusion of the study. GraphPad Prism 8.0 software (RRID: SCR_002798) was used for statistical analysis and graphing. Bar graphs show means with standard error of the mean and symbols for individual mouse values. Two-way ANOVA was used to determine statistically significant differences when comparing by genotype and injury between groups. Repeated measures two-way ANOVA (RM ANOVA) was used to assess differences between genotypes over time in the longitudinal MRI analysis and within behavioral assessments. Corrections for multiple comparisons were done using Sidak’s test. An alpha-level of p value < 0.05 was considered statistically significant.

## RESULTS

### *Sarm1* deletion protects against CC atrophy and axon-myelin pathology in chronic TBI

The potential for *Sarm1* inactivation to provide long term axon protection and/or to mitigate post-traumatic neurodegeneration was first evaluated using transmission EM (Figure 1). Before thin sectioning for EM, sagittal brain slices were osmium-stained to label myelin and examined with bright field microscopy to measure the CC width, which is a clinically relevant measure of white matter pathology in TBI and neurological diseases [19]. Both *Sarm1* WT and *Sarm1* KO mice developed significant CC atrophy by 10 weeks after TBI, yet the extent of atrophy was significantly less in the *Sarm1* KO mice (Figure 1A). For in-depth analysis of the underlying pathology associated with changes in CC size, the tissue slices were then thin sectioned and imaged by transmission EM at subcellular resolution to examine individual myelinated axons within the same region of the CC. This quantification revealed a significant reduction of intact axons due to TBI in the *Sarm1* WT mice that was rescued by *Sarm1* deletion (Figure 1B).

**Figure 1.**
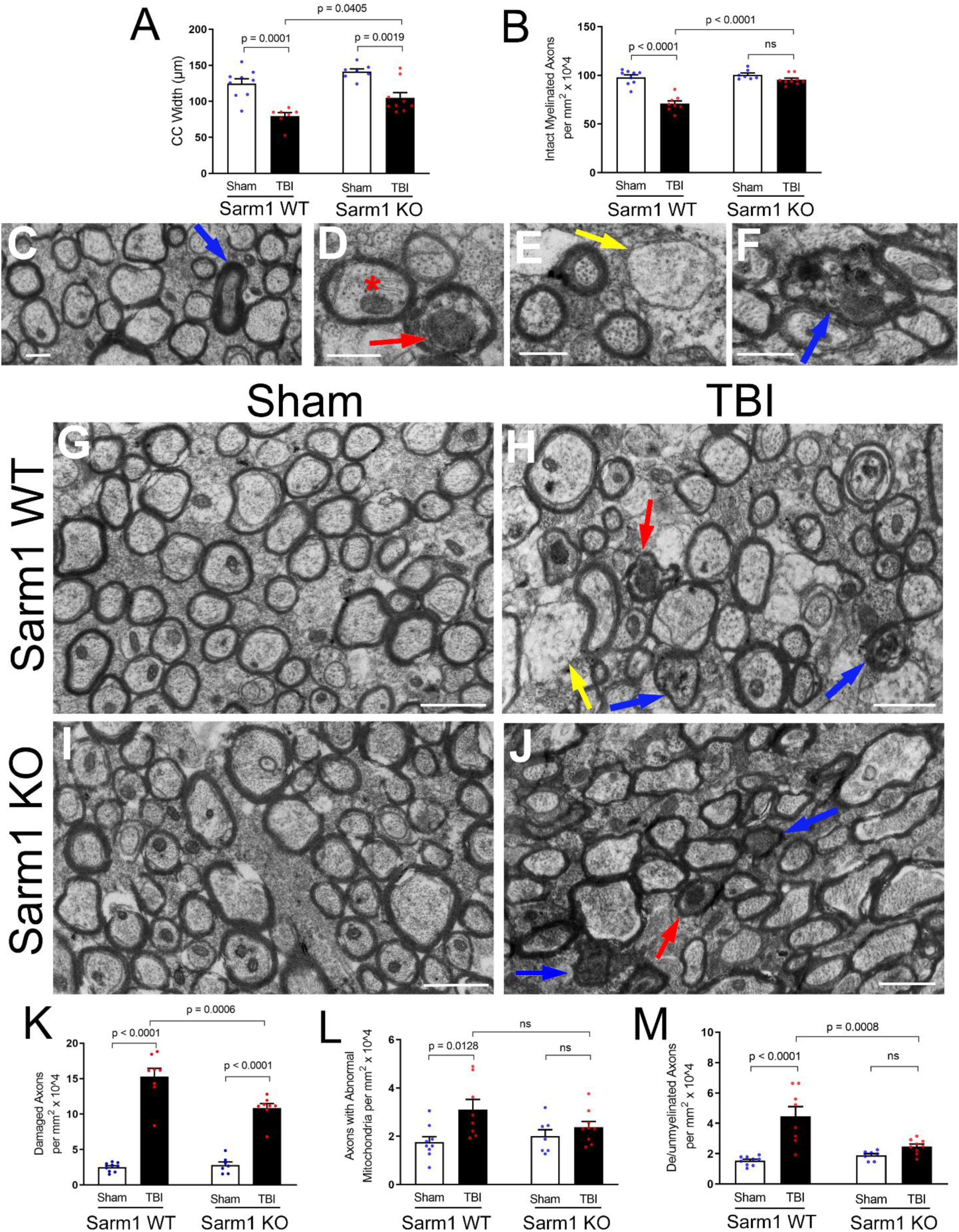
*Sarm1* deletion reduces corpus callosum atrophy and axon pathology at 10 weeks post-TBI. **A:** *Sarm1* deletion attenuates CC atrophy that develops after TBI. **B:** *Sarm1* deletion preserves intact, myelinated axons after TBI**. C-F:** Representative images of pathological features from high resolution electron microscopy in CC sagittal sections. **C:** Example from a *Sarm1* WT sham mouse to show the cytoskeleton, mitochondria and surrounding myelin sheath of intact axons in contrast with a rare damaged axon with a densely compacted cytoskeleton (blue arrow). **D:** *Sarm1* WT TBI mouse example of an axon with an abnormally large mitochondrion (red arrow) and an axon with a typical mitochondrion (red asterisk). Mitochondria were considered abnormal when swollen to > 50% of the axon area. **E:** Demyelinated axon (yellow arrow) lacking ensheathing myelin but with otherwise intact cytoskeleton in a *Sarm1* WT TBI mouse. **F:** Damaged axon (blue arrow) with vesicle accumulation and cytoskeletal breakdown in a *Sarm1* KO TBI mouse. **G-J:** Representative electron microscopy images from CC sagittal sections for *Sarm1* WT sham (G) and TBI (H) mice in comparison with *Sarm1* KO sham (I) and TBI (J) mice. Arrows identify examples of damaged axons with accumulated vesicles and/or compacted cytoskeleton (blue), abnormal mitochondria (red), or demyelination (yellow). **K-M:** Quantification of axon and myelin pathology at 10 weeks after TBI or sham procedure. *Sarm1* deletion reduces chronic stage axon damage (K), normalizes mitochondria morphology (L), and eliminates TBI-induced demyelinated component of de/unmyelinated axons (M). *Sarm1* WT: n = 9 sham, n =8 TBI. *Sarm1* KO: n = 7 sham, n = 9 TBI. ns = not significant. C-F, scale bars = 0.5 µm. G-J, scale bars = 1 µm.

Myelinated axons exhibit subcellular features that can distinguish intact axons from indicators of axon damage including swollen mitochondria, compacted cytoskeletal structure, accumulation of vesicles or debris, or loss of ensheathing myelin (Figure 1C-F). After the sham procedure, *Sarm1* WT mice illustrate the healthy adult high density of myelinated axons in this anterior region of the CC (Figure 1G). As previously characterized in this concussive TBI model [51, 58], TBI results in damaged axons that are dispersed among adjacent intact axons and may also exhibit demyelination in *Sarm1* WT mice (Figure 1H). *Sarm1* KO mice exhibit normal appearing myelinated axons after the sham procedure and dispersed damaged axons after TBI (Figure 1I, J). Importantly, while both *Sarm1* WT and *Sarm1* KO mice exhibited axon damage after TBI, *Sarm1* deletion significantly reduced the frequency of damaged axons (Figure 1K). *Sarm1* deletion also normalized the frequency of TBI-induced mitochondrial pathology and demyelination (Figures 1L, M).

### Longitudinal *in vivo* MRI detects CC atrophy after TBI and attenuation by *Sarm1* deletion

Longitudinal MRI studies were conducted to advance the translational impact while further evaluating the effects of *Sarm1* deletion on white matter integrity and CC atrophy. Each mouse was scanned prior to injury (i.e., baseline), and with follow up scans at acute and chronic stages after TBI. While DTI measures of white matter integrity have long been effectively used for analysis of adult mouse CC [75, 78], volume measurements have only recently been validated for quantification of atrophy and hypertrophy of the relatively small structures of adult mouse brains [3, 9]; Therefore, an initial cross-sectional *in vivo* study was conducted in C57BL/6 mice to optimize the MRI outcome measures. DTI analysis at 10 weeks after sham or TBI demonstrated significant changes after TBI in CC integrity while high resolution T2-w volumetrics detected significant CC atrophy that was validated by post-imaging neuropathology (Figure S2, Online Resource 1).

Longitudinal MRI studies were then conducted to compare *Sarm1* WT and *Sarm1* KO mice. Volumetric analysis showed significant atrophy of the CC between baseline and 10 weeks in *Sarm1* WT mice (Figure 2A-C) that was attenuated in *Sarm1* KO (Figure 2D-F). Across time points, DTI detected reduced white matter integrity after TBI based on reduced fractional anisotropy (FA) in both *Sarm1* WT mice (Figure 2G, I) and *Sarm1* KO mice (Figure 2H, I). Progressive decrease in FA was driven by decreased axial diffusivity (AD) at 3 days post-TBI (Figure 2J) and subsequent elevation of radial diffusivity (RD) at 10 weeks post-TBI (Figure 2K). *Sarm1* genotype resulted in a significant effect of higher AD values in *Sarm1* KO mice (Figure 2J).

**Figure 2.**
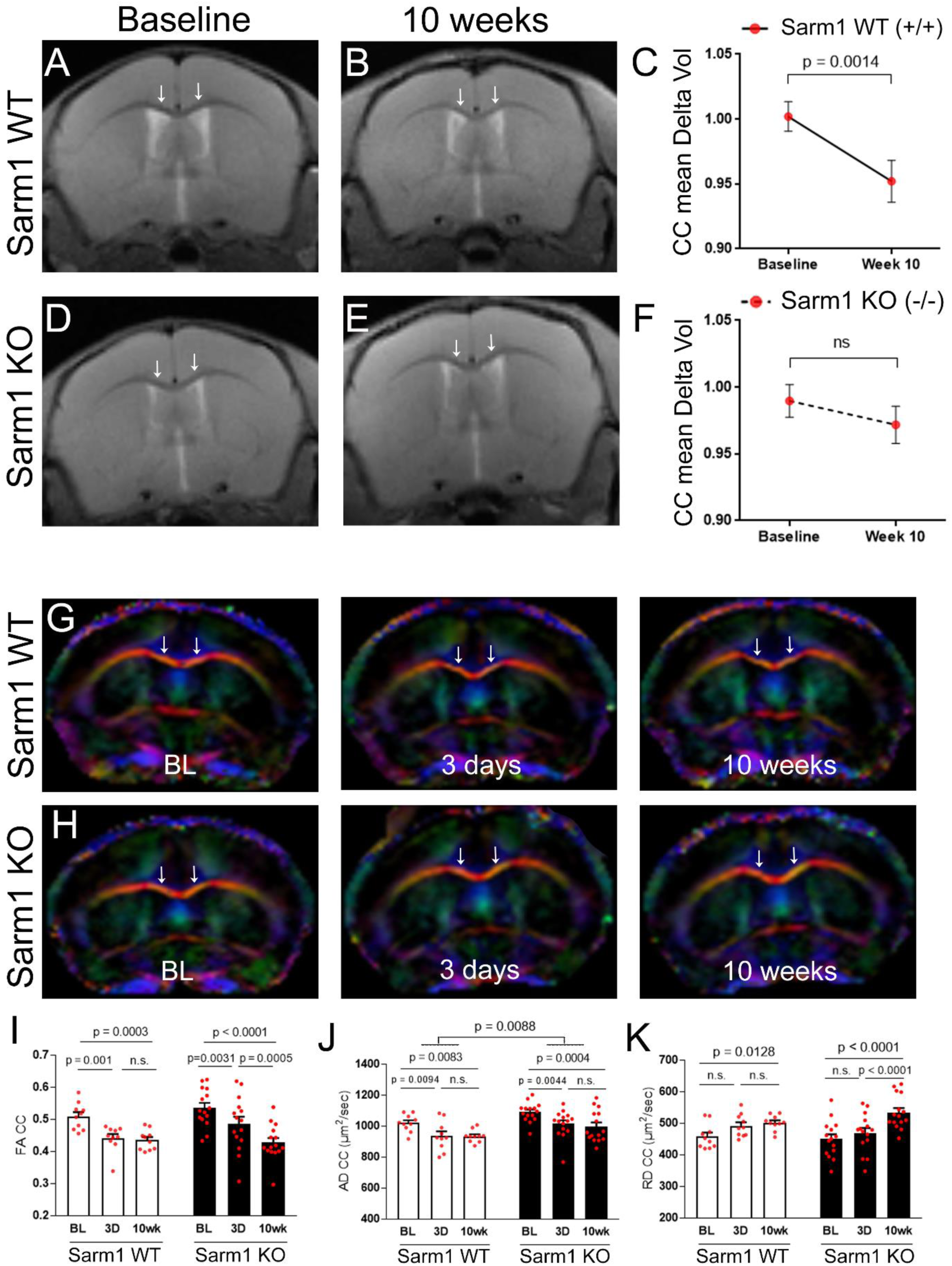
Progression of corpus callosum atrophy after TBI is sufficient for MRI detection in live mice and is attenuated by *Sarm1* deletion. **A-C:** TBI induces significant CC atrophy in *Sarm1* WT mice. T2-weighted images showing the coronal view at the level of the impact site at baseline (A, before surgery) and at 10 weeks (B) post-TBI/sham procedures. Quantification of volume change in CC regions under the impact site (C). **D-F:** In *Sarm1* KO mice, CC atrophy is not detected in representative T2-weighted images (D, E) or based on changes in CC volume (F). **G-H:** Direction encoded color images of diffusion tensor imaging (DTI) from a longitudinal MRI series at baseline (BL) and again at 3 days and 10 weeks post-TBI or sham procedures. Colors represent fiber directions as red (medial-lateral), blue (anterior-posterior), and green (superior-inferior). **I-K:** Quantification of DTI measures reveals a chronic progression of CC pathology following TBI. Fractional anisotropy (FA) significantly decreases over time following TBI (I). The acute change is driven by a decrease in axial diffusivity (AD) between baseline and 3 days (J). *Sarm1* KO mice have significantly higher AD values than *Sarm1* WT mice (J). The FA at 10 weeks corresponds with a delayed increase in radial diffusivity (RD) in *Sarm1* KO mice (K). Arrows indicate medial CC regions. *Sarm1* WT: n = 10 TBI; *Sarm1* KO: n = 15 TBI. ns = not significant.

### Immunohistochemistry shows reduced neuroinflammation in addition to with reduced CC atrophy and myelin loss after TBI in mice with *Sarm1* deletion

Microglia and astrocyte responses have been linked to chronic white matter damage and atrophy [40, 44, 45, 49, 58, 79]. Therefore, immunohistochemistry was used to evaluate CC atrophy and myelination (Figure 3A-D) and examine CC neuroinflammation in adjacent sections (Figure 4). CC width was measured in coronal sections with myelin immunolabeled for MOG and cytoarchitecture was evaluated using DAPI to stain nuclei (Figures 3E and S2, Online Resource 1). CC width was reduced at 10 weeks after TBI in comparison with sham mice (Figure 3E). Quantification of the pixel area of MOG immunoreactivity within the CC showed loss of myelin (Figure 3F), in agreement with the distributed demyelination observed by EM (Figure 1M). *Sarm1* KO mice had significantly less CC atrophy and less myelin loss after TBI as compared to *Sarm1* WT mice (Figure 3E, F).

**Figure 3.**
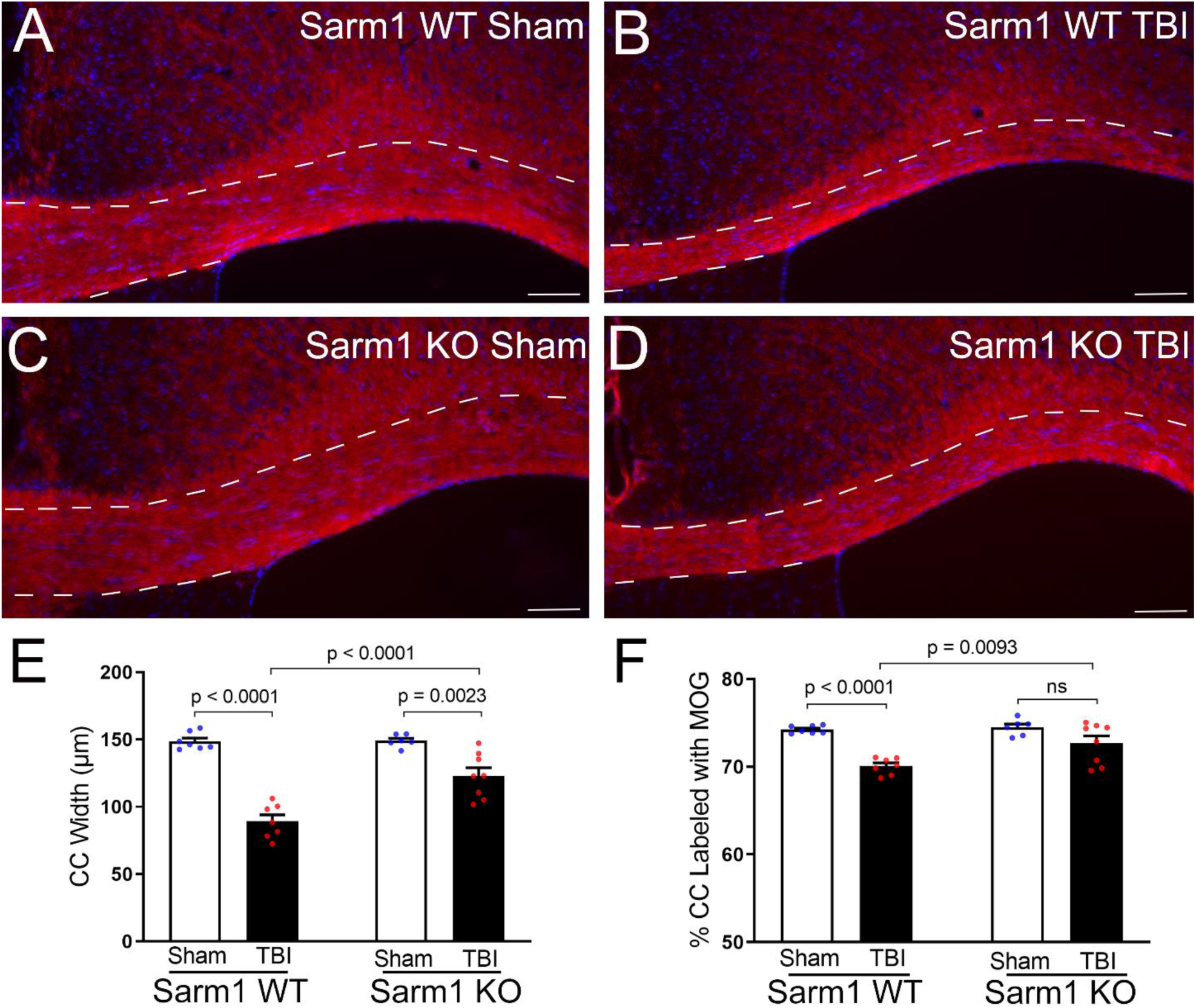
*Sarm1* deletion reduces corpus callosum atrophy and myelin loss at 10 weeks post-TBI. **A-D:** Representative images from CC coronal sections of *Sarm1* WT and *Sarm1* KO mice after sham or TBI procedures. Myelin is detected with immunolabeling for MOG (red). DAPI nuclear stain is shown in blue. CC borders are indicated by dashed lines. **E:** *Sarm1* deletion attenuates CC atrophy, which is quantified based on the CC width. **F:** TBI results in significant myelin loss as detected by reduced MOG immunoreactivity. Myelin loss after TBI is significantly reduced in *Sarm1* KO mice compared to *Sarm1* WT mice. *Sarm1* WT: n = 7 sham, n = 7 TBI. *Sarm1* KO: n = 6 sham, n = 8 TBI. ns = not significant. A-D, scale bars = 100 µm.

**Figure 4.**
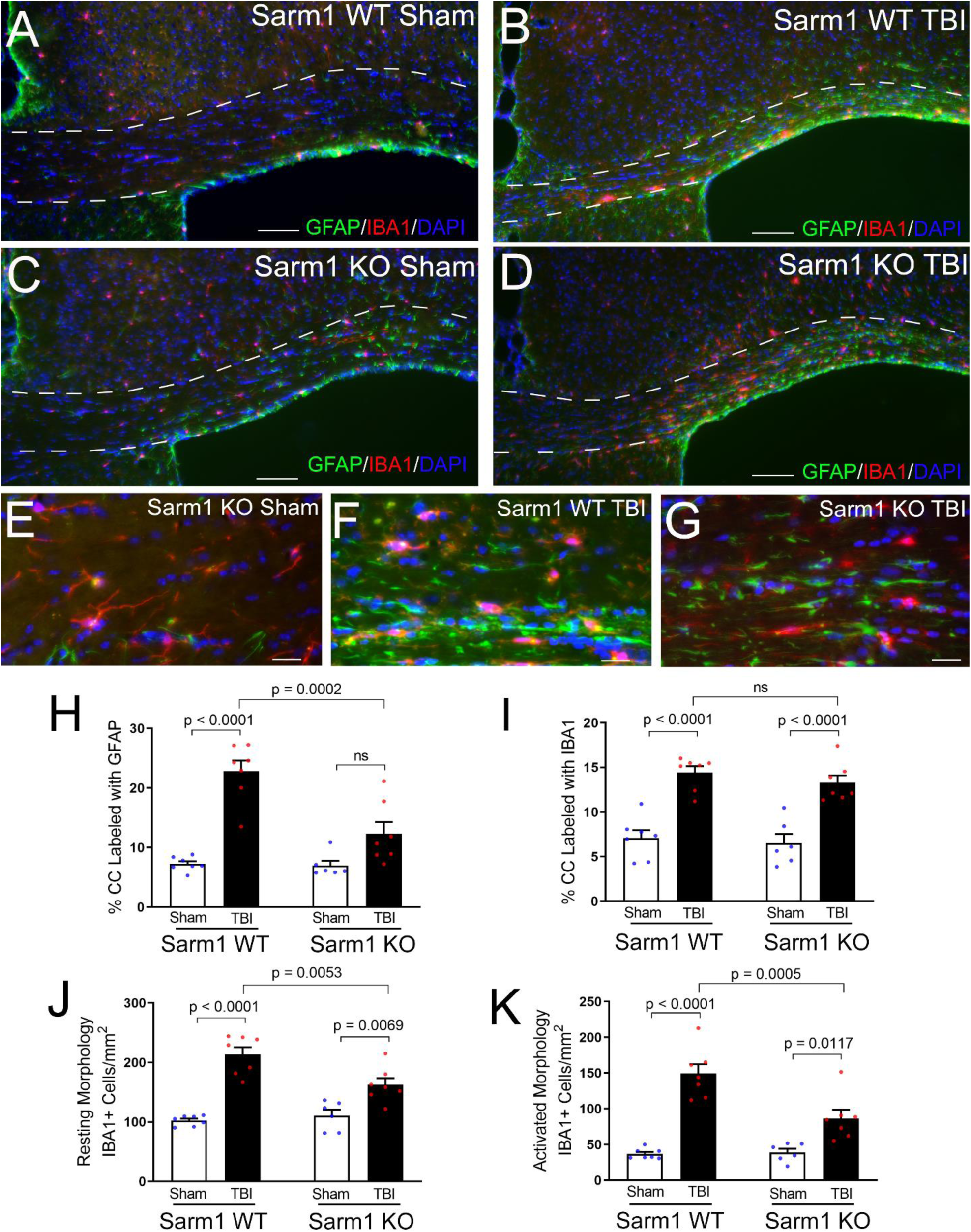
*Sarm1* deletion reduces neuroinflammation at 10 weeks post-TBI. **A-D:** Representative images from CC coronal sections of *Sarm1* WT (A, B) and *Sarm1* KO (C, D) mice after sham (A, C) or TBI (B, D) procedures. Neuroinflammation is detected with markers of astrocytes (GFAP, green) and microglia (IBA1, red). DAPI nuclear stain shown in blue. The CC borders are indicated by dashed lines. **E-G**: Higher magnification examples of astrocyte (GFAP) and microglia (IBA1) morphology. In sham mice (E), astrocytes and microglia exhibit homeostatic morphology with thin processes. Following TBI (F, G), reactive astrocytes and microglia have intensely immunolabeled cell bodies and shorter, thicker, processes. **H:** *Sarm1* deletion significantly reduced astrogliosis after TBI, based on GFAP immunolabeling within the CC area. **I:** The microglial response also indicated CC neuroinflammation after TBI, based on IBA1 immunolabeling, but did not detect differences due to *Sarm1* deletion. **J-K:** More detailed counting of IBA1 immunolabeled (+) cells revealed that *Sarm1* deletion significantly reduced the frequency of both resting (J) and activated (K) microglia after TBI. *Sarm1* WT: n = 7 sham, n = 7 TBI. *Sarm1* KO: n = 6 sham, n = 7 TBI. ns = not significant. A-D, scale bars = 100 µm. E-G, scale bars = 25 µm.

Neuroinflammation was estimated by immunolabeling for reactive astrocytes and microglia using GFAP and IBA1, respectively (Figure 4). TBI resulted in more intense immunoreactivity of individual cells and an increase of overall immunolabeling for both GFAP and IBA1 (Figure 4A-G). Immunolabeling within the CC after TBI was reduced in *Sarm1* KO mice for GFAP (Figure 4H) but not for IBA1 (Figure 4I). Further analysis to count the IBA1 immunolabeled cells within the CC revealed that *Sarm1* deletion reduced microglia with either a resting state morphology (Figure 4J) or an activated morphology (Figure 4K) after TBI. Taken together, these results show that the reduction in CC atrophy and myelin loss seen in *Sarm1* KO mice after TBI was accompanied by a reduced neuroinflammatory response.

### *Sarm1* deletion improves functional outcome measures in chronic stage TBI

Behavioral assessments targeting CC axons were selected to test whether the beneficial effects of *Sarm1* deletion on TBI pathology translated to improved functional outcome measures. This TBI model did not cause overt symptoms at any time out through 10 weeks post-TBI. Therefore, an initial set of experiments in C57BL/6J mice evaluated two assays associated with CC axon-myelin pathology to determine whether either assay revealed deficits during this late phase of TBI. Miss-step wheel running is a motor skill task that engages CC axons and is sensitive to myelination status [35, 57, 78]. C57BL/6 mice showed a deficit in learning to run on the Miss-step wheels after TBI as compared to sham mice (Figure S3, Online Resource 1). C57BL/6 mice with CC pathology from experimental demyelination or repetitive mild exhibit social interaction deficits TBI [59, 91]. However, with the current single impact concussive model of TBI, C57BL/6 mice did not show social interaction deficits (Figure S4, Online Resource 1). Based on the results of these assays, the Miss-step wheel task was selected for functional assessment of TBI deficits in *Sarm1* WT versus *Sarm1* KO mice.

Motor learning and performance on the Miss-step wheels was assessed in *Sarm1* WT and *Sarm1* KO mice from 8-10 weeks post-TBI. *Sarm1* KO mice ran at a faster average velocity compared to *Sarm1* WT mice during the learning phase of the assay although this improvement did not reach statistical significance (p = 0.0821) (Figure 5A). Further analysis of running parameters showed that *Sarm1* KO mice ran on the wheels more times during the learning phase compared to *Sarm1* WT mice (Figure 5B). Additionally, the cumulative distance traveled by *Sarm1* KO mice during the learning phase is further for *Sarm1* KO mice than for *Sarm1* WT mice (Figure 5C).

**Figure 5.**
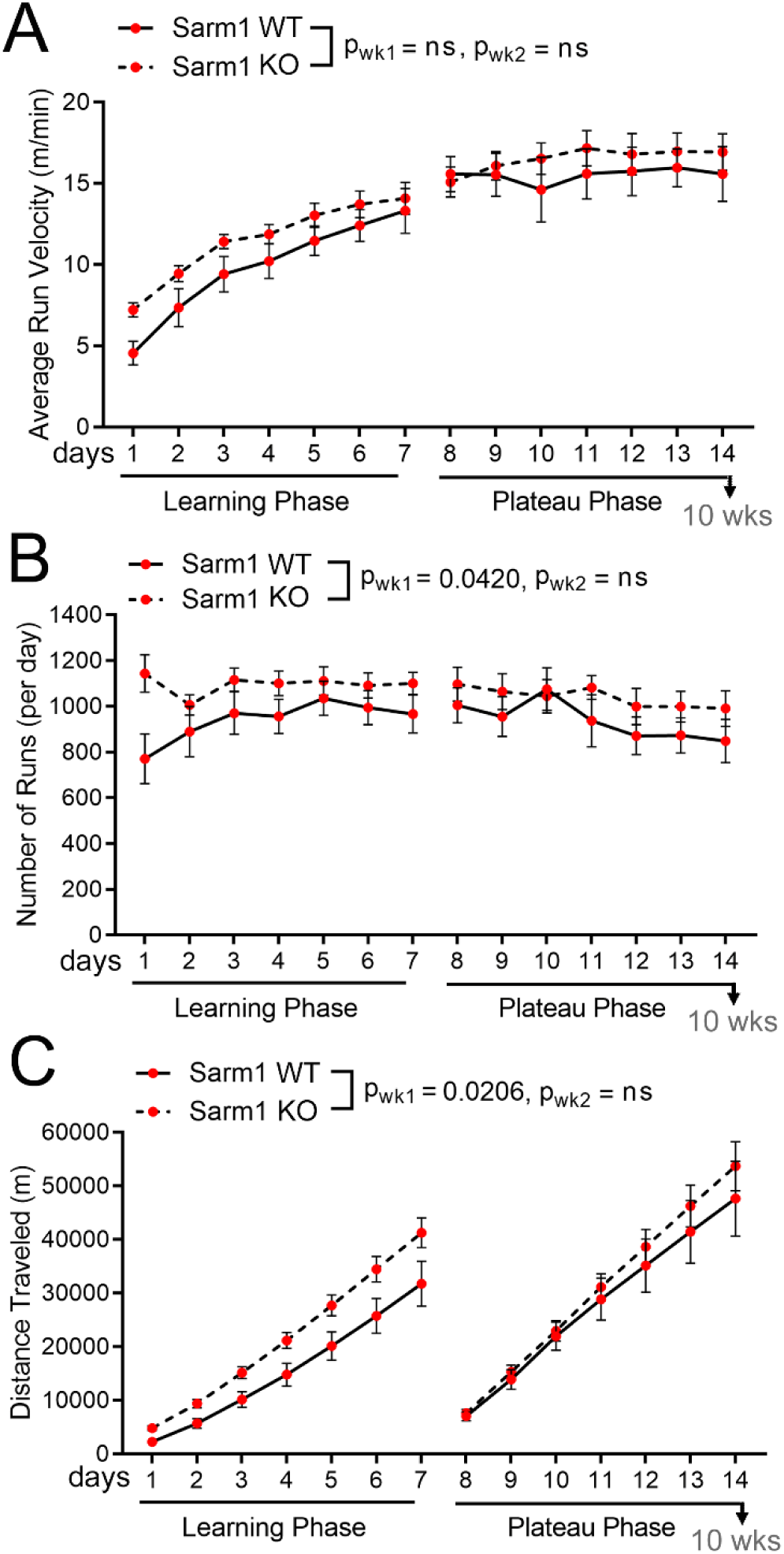
*Sarm1* deletion improves motor learning in chronic stage TBI. Miss-step wheels have irregularly spaced rungs to assess motor skill learning (week 1) followed by a plateau phase (week 2) that tests bilateral sensorimotor function. **A:** *Sarm1* KO mice more quickly learn to run at a faster average velocity compared to *Sarm1* WT mice, but this improvement does not reach statistical significance (p = 0.0821). **B:** *Sarm1* KO mice run on the wheels more frequently than *Sarm1* WT during the learning phase. **C:** Running behavior accumulates to increased total distance traveled during the learning phase in *Sarm1* KO mice compared to *Sarm1* WT mice. There were no statistically significant differences between genotypes in the three measures during the plateau phase (A-C). *Sarm1* WT: n = 9 TBI. *Sarm1* KO: n = 15 TBI. ns = not significant.

Sleep behavior was selected in place of social interaction as a highly translational assessment for post-traumatic neurodegeneration (Figure 6). Sleep disorders are common in patients with chronic TBI and sleep patterns may contribute to neurodegeneration, particularly white matter degeneration [2, 63, 72]. Sleep data was analyzed for *Sarm1* WT and *Sarm1* KO mice during two light/dark cycles. In *Sarm1* WT mice, TBI causes an overall difference from shams in the percent of time spent sleeping (Figure 6A). This difference was not found in *Sarm1* KO mice (Figure 6B). More specifically, *Sarm1* WT mice slept less during the lighted period that is the sleep phase for nocturnal mice (Figure 6C) without a change during the awake phase (Figure 6D). *Sarm1* KO mice did not exhibit a sleep deficit after TBI, as compared to shams (Figure 6C, D).

**Figure 6.**
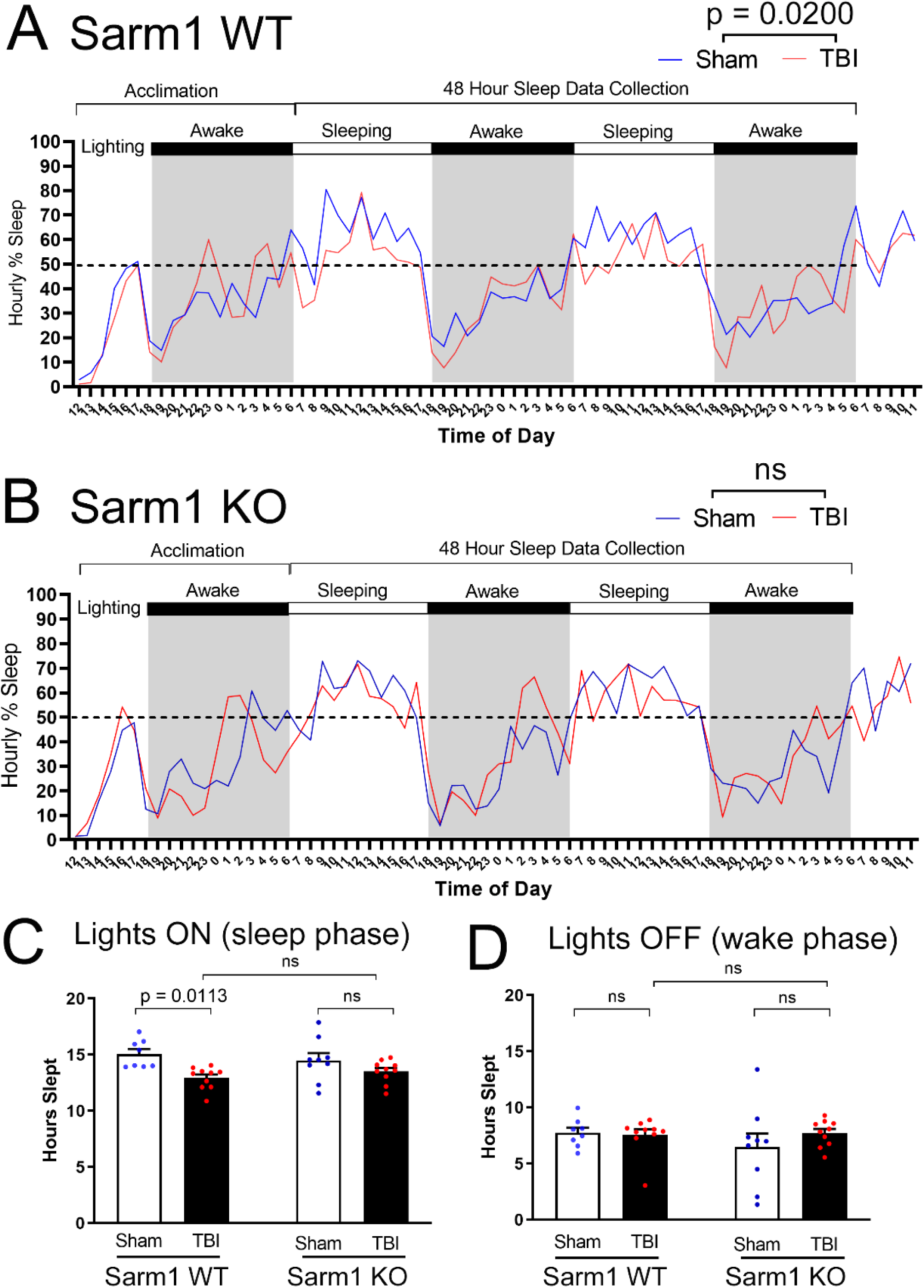
*Sarm1* deletion normalizes time spent sleeping in the chronic stage of TBI: Mice were single housed in cages equipped with a PiezoSleep Mouse Behavioral Tracking System for 72 hrs during the eighth week after TBI. **A-B:** Mice were acclimated for 18 hrs. Statistical analysis focused on data collected during 48 hrs across two complete cycles of lights on (white bars) and lights off (dark bars and gray background). Recordings continued for a subsequent 6 hrs to show the final wake/sleep transition. In *Sarm1* WT mice (A), the time spent sleeping per hour is significantly different between sham and TBI, with the injured mice appearing to sleep less during the normal sleeping period when lights are on. In *Sarm1* KO mice (B) the sleep pattern was not different between sham and TBI conditions during the 48 hr data collection period. **C:** TBI significantly reduced the time spent sleeping for *Sarm1* WT mice during the normal sleeping period when lights are on. *Sarm1* KO mice did not exhibit sleep loss after TBI. **D:** During the dark period, sleep time was not different based on injury or genotype. *Sarm1* WT: n = 8 sham, n = 10 TBI. *Sarm1* KO: n = 9 sham, n = 10 TBI. ns = not significant.

## DISCUSSION

The combined results from this study support a direct role of axonal injury in post-traumatic neurodegeneration and provide key pre-clinical evidence for TBI as a clinical indication for treatments to inhibit SARM1. Genetic inactivation of the gene for SARM1, which mediates a conserved axon degeneration pathway, had a beneficial effect on chronic white matter injury after TBI. Analysis of CC atrophy in longitudinal and cross-sectional MRI studies was an effective outcome measure in mice that is a clinically useful measure of post-traumatic neurodegeneration. Complementary neuropathological techniques validated CC atrophy measures and identified beneficial effects from deletion of the *Sarm1* gene on chronic stage axon damage, demyelination, and neuroinflammation. Importantly, *Sarm1* deletion preserved healthy axons and prevented the progression of CC atrophy after TBI. *Sarm1* deletion also ameliorated functional deficits which, together with the pathological and radiological benefits, indicate improved outcome trajectory after TBI.

Insights afforded by the subcellular detail of EM (Figure 1) address important gaps in the interpretation of long-term effects of *Sarm1* deletion in experimental TBI. In the TBI model used in the current studies, EM data from the acute time point of 3 days [51] and chronic phase of 10 weeks post-TBI (Figure 1) are matched by scan time points in our longitudinal MRI of live mice (Figure 2). Prior studies in *Sarm1* KO mice have shown significant axon protection in the acute (2 hours - 3 days) or chronic (2-6 months) phase after closed head TBI based on β-amyloid precursor protein (β-APP) immunohistochemistry [34, 56] or using Thy1-YFP fluorescence to visualize axonal varicosities [51, 92]. The observation of subcellular details of axon and myelin ultrastructure afforded by EM informs a fuller interpretation of myelinated axon health.

EM enables quantification of abnormal swollen mitochondria that are an early feature of axonal injury linked to Wallerian degeneration [86, 88] and have important implications for the SARM1 pathway [36, 47]. Mitochondrial dysfunction challenges axon energy metabolism and creates a vulnerable axon state [36, 65]. Mitochondrial stress, trauma or other insults lead to loss of nicotinamide mononucleotide adenylyl-transferase 2 (NMNAT2), an NAD-synthesizing enzyme, in axons that depletes NAD+ and raises the level of its precursor nicotinamide mononucleotide (NMN) [20, 47]. Low NAD+ and high NMN levels regulate release of auto-inhibition of SARM1 NADase activity, which further depletes NAD+ and axon energy stores [11, 21, 38, 73, 76]. Axons with swollen mitochondria were significantly increased at 3 days after TBI in *Sarm1* WT and *Sarm1* KO mice [51]. Compared to this acute data, axons with swollen mitochondria were less frequent at 10 weeks but still significantly increased after TBI in *Sarm1* WT mice (Figure 1), which indicates the potential for continued activation of SARM1.

EM identifies damaged axons based on breakdown and compaction of the cytoskeleton and/or accumulated vesicles due to impaired axonal transport (Figure 1). The frequency of damaged axons and the effect of *Sarm1* deletion is very similar at the 10 week time point (Figure 1) as compared to 3 days post-TBI [51]. The slow clearance of degenerating axons in the CNS may play a part in this result [85]. Alternatively, surviving axons may succumb to insults in a late phase after TBI and *Sarm1* deletion may have on ongoing benefit across acute and chronic time points. For example, axons continue to initiate Wallerian degeneration long after stretch injury that models TBI forces [54]. Clinically, persistent neuroinflammation in white matter is associated with axon damage in the CC in human postmortem cases several years after TBI [39].

Finally, EM is the gold standard to quantify demyelination, i.e. loss of the myelin sheath around otherwise healthy axons. TBI induced demyelination in *Sarm1* WT mice that was not found in *Sarm1* KO mice at 10 weeks post-TBI or sham procedures (Figure 1). *Sarm1* is expressed mainly in neurons but a low level of expression in oligodendrocyte lineage cells could have a role in the lack of TBI-induced demyelination in *Sarm1* KO mice [43]. However, the effect of *Sarm1* deletion may not be an autonomous effect in myelinating oligodendrocytes as zebrafish studies have shown a glioprotective effect of *Sarm1* deletion is dependent on axon protection [82]. Loss of myelin along otherwise healthy axons can slow information processing speed and desynchronize neural circuits; myelin also protects from insults and provides trophic support to axons [6]. Therefore, demyelination that is resolved by *Sarm1* deletion may have ongoing benefit at 10 weeks post-TBI that can impact axon health and neural circuit function.

The current study is the first to use translational measures of MRI with CC atrophy to evaluate the effects of *Sarm1* deletion after TBI. Based on histological measures of CC width, our prior studies in this TBI model did not detect significant CC atrophy at the 3 day, 2 week, or 6 week time points [51, 58]. In contrast, significant CC atrophy was evident by histological measures at the longer time point of 8 weeks post-TBI [51, 52] and now at 10 weeks post-TBI (Figures 1A, 3E and S2, Online Resource 1). At this 10 week time point, these histological measures validate the changes in CC volume in live mice determined by quantitative MRI (Figure 2C and S2, Online Resource 1). Significantly, the histological and MRI measures corroborate attenuation of CC atrophy in *Sarm1* KO mice as compared to *Sarm1* WT mice (Figures 1A, 2F, 3E). These longitudinal MRI studies also analyzed DTI FA as a measure of CC microstructure [87, 90]. Reduced FA values indicated an injury effect at 3 days and at 10 weeks after TBI, as compared to baseline (Figure 2I-K and S2, Online Resource 1). A main effect of *Sarm1* genotype was only observed for the AD parameter (Figure 2J) that is most often associated with axonal anisotropy within white matter voxels. Interpreting reduced AD as an indicator of axon pathology and increased RD as an indicator of myelin pathology can be useful when each pathology predominates [75, 89]. The presence of simultaneous pathology of axon damage and demyelination is more difficult to detect and interpret using DTI, and is further complicated in the presence of significant edema or neuroinflammation [87, 89]. Histological measures confirm the presence of simultaneous axon damage, demyelination, and neuroinflammation at 10 weeks after TBI along with significant reduction of each pathology with *Sarm1* deletion (Figures 1, 3, 4), which is not fully appreciated using the DTI parameters (Figure 2I-K).

Pathological and structural benefits of *Sarm1* deletion were complemented by behavioral studies that show amelioration of functional deficits (Figures 5, 6). Our results extend beyond previous reports in *Sarm1* KO mice to now examine complex behaviors during the chronic phase post-TBI. Studies in a weight drop model of TBI used the neurological severity score battery of simple tasks from 2 hours through 4 weeks post-TBI and found reduced deficits in *Sarm1* KO versus *Sarm1* WT mice only during the first week [34]. Studies of repetitive TBI showed normalization toward sham responses in *Sarm1* KO for motor performance and memory deficits during the first week and context fear discrimination at 4 weeks [56]. With assessments conducted beyond 8 weeks post-TBI, *Sarm1* deletion reduced deficits during motor learning (Figure 5) and normalized the time spent sleeping (Figure 6). The Miss-step wheel running system (Figure 5) and the piezoelectric sleep system (Figure 6) use automated multi-day continuous data collection of spontaneous complex behaviors, which may be advantageous for analysis of subtle deficits after TBI.

Limitations of the experimental design should be considered in the interpretation of the results. The *Sarm1* KO mice have been backcrossed to the C57BL/6 strain but may harbor genes associated with the embryonic stem cell origin in the 129 background strain [84]. However, the findings regarding axon degeneration in this line of *Sarm1* KO mice have been confirmed in additional *Sarm1* KO mice generated using CRISPR technology [84]. In addition, all experiments in the current study used *Sarm1* WT and *Sarm1* KO littermates to minimize the variability due to genetic background. The assessment of sleep disorders after TBI used a non-invasive screening approach to avoid surgical procedures and electrode placement through skull burr holes that would be required for electrophysiology. The sleep behavior in the *Sarm1* KO mice appears normalized to the sham sleep pattern but electrophysiological recordings would be needed for a more in-depth comparison of sleep architecture. Finally, the neuropathological and radiological analyses focused on white matter, and specifically the CC. Further studies would be of interest to better understand the relationship of axon damage and the progression of white matter pathology relative to broader neural circuits and gray matter pathology, including analysis of synapse loss that is associated with chronic neuroinflammation after TBI [1].

## CONCLUSIONS

These results demonstrate that genetic inactivation of *Sarm1* improves the outcome trajectory after TBI based on pathological, radiological, and functional measures. These studies advance strategies to develop TBI treatments for axon damage by demonstrating a genetic proof-of-concept of the long-term benefit of *Sarm1* deletion. The therapeutic potential of SARM1 inhibitors has drawn intense interest and a small molecule inhibitor of SARM1 has already been developed that recapitulates *in vitro* aspects of the *Sarm1* KO phenotype [36, 48, 74]. The current studies also highlight CC atrophy as an important outcome measure of white matter degeneration after TBI in mice that may have translational relevance as a biomarker for clinical studies [28]. White matter degeneration may be a tractable therapeutic target for TBI and chronic traumatic encephalopathy, with potential application to other neurodegenerative diseases including Alzheimer’s disease and multiple sclerosis [4, 14, 16, 70].

## Acknowledgements

The authors thank Dr. Christina Marion and Dr. Krystal Schaar Valenzuela for technical advice. We appreciate the support of the Center for Neuroscience and Regenerative Medicine Preclinical Models Core, Translational Imaging Core, and Translational Therapeutics Core and the Biomedical Instrumentation Center at the Uniformed Services University. These studies were funded by the U.S. Department of Defense and the Uniformed Services University through the UCSF-USUHS Partnership: Brain Injury and Disease Prevention, Treatment, and Research and the Center for Neuroscience and Regenerative Medicine. The authors declare no competing financial interests. Opinions are those of the authors and do not represent the University, the Department of Defense, or the federal government.

---

**Supplemental Figure S1:**
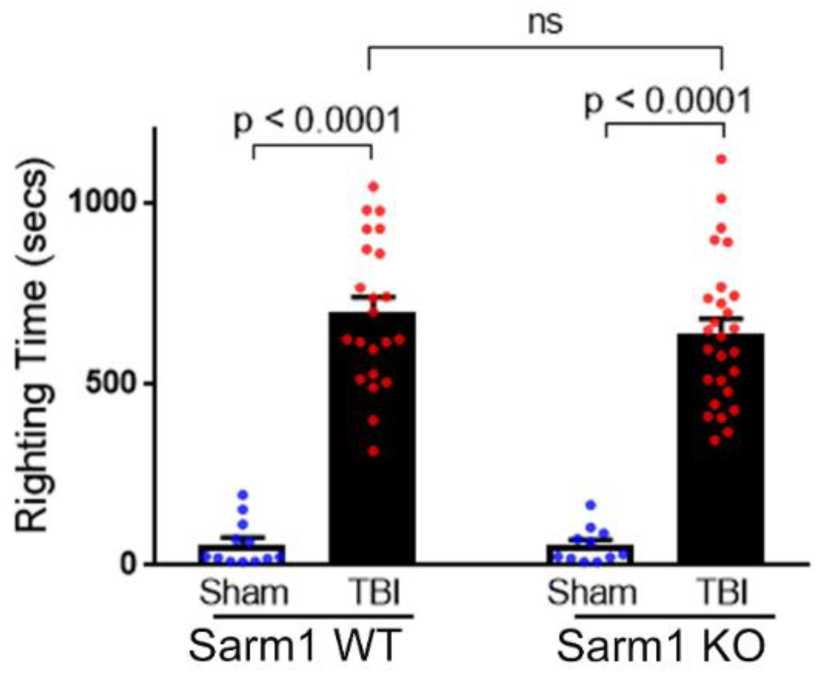
Righting reflex times after sham or TBI surgical procedure in *Sarm1* WT and *Sarm1* KO mice. Post-surgical righting reflex times were measured as the time interval from the end of anesthesia until mice returned to the upright position. Righting time is shown for all mice used in experiments. Prolonged righting reflex times are an indicator of TBI severity that serve as a measure of arousal that estimates loss of consciousness after TBI or sham procedures. Righting reflex time did not differ between male and female littermates, so the sexes were combined for analysis by genotype. Righting reflex time was significantly prolonged after TBI as compared to the sham procedure for *Sarm1* WT (p < 0.0001) and *Sarm1* KO (p <0.001) mice without a difference based on genotype (p = 0.8287; Two-way ANOVA; F(1,68) = 183).

**Supplemental Figure S2.**
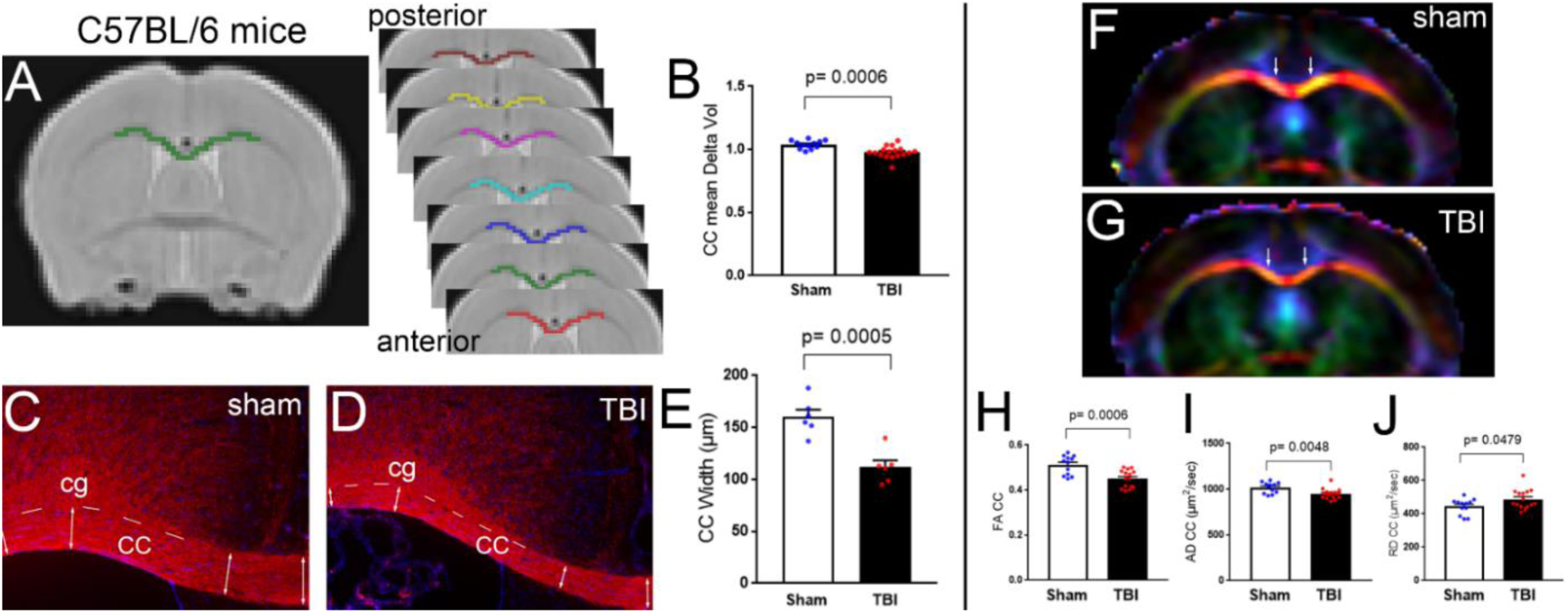
TBI reduces white matter integrity and produces significant corpus callosum (CC) atrophy in C57BL/6 mice. A cross-sectional *in vivo* MRI study used male C57BL/6J mice between 9-10 weeks post-TBI/sham. After scanning, mice were perfused for tissue analysis at 10 weeks post-TBI/sham. **A-B:** CC volume changes were calculated as the change in volume compared to a registered template image of all animals in the study. Seven ROIs were manually drawn on coronal images (125 µm thickness) encompassing the rostro-caudal CC over the lateral ventricle and underneath the impact site at bregma (A). TBI significantly reduced the mean CC volume, as compared to the sham procedure (B) **C-E:** Post-imaging neuropathology validated CC atrophy after TBI. Myelinated fibers were immunolabeled for myelin oligodendrocyte glycoprotein (MOG; red) and cellular distribution was detected using nuclear counterstain (DAPI; blue) (C, D). The CC borders (dashed lines) were evident by the pattern of myelinated fibers oriented in the medial-lateral direction in the CC in contrast to the myelinated fibers of the cingulum (Cg) that align in the rostro-caudal (i.e. anterior-posterior) direction. Double headed arrows (C, D) show examples of sites for CC width measurement for quantification (E). **F-J:** MRI diffusion tensor imaging (DTI) illustrates fractional anisotropy signal in the CC (white arrows) and adjacent regions in direction encoded color maps (F, G). Colors represent fiber directions as red (medial-lateral), blue (anterior-posterior), and green (superior-inferior). TBI significantly reduced the CC fractional anisotropy (FA; H) which was driven by reduced axial diffusivity (AD; I) and increased radial diffusivity (RD; J). MRI n = 12 per condition; male C57BL/6J mice (RRID:IMSR_JAX:000664), Jackson Laboratories, Bar Harbor, ME). Neuropathology n = 6 per condition randomly selected from the MRI cohort. Student’s *t*-test.

**Supplemental Figure S3.**
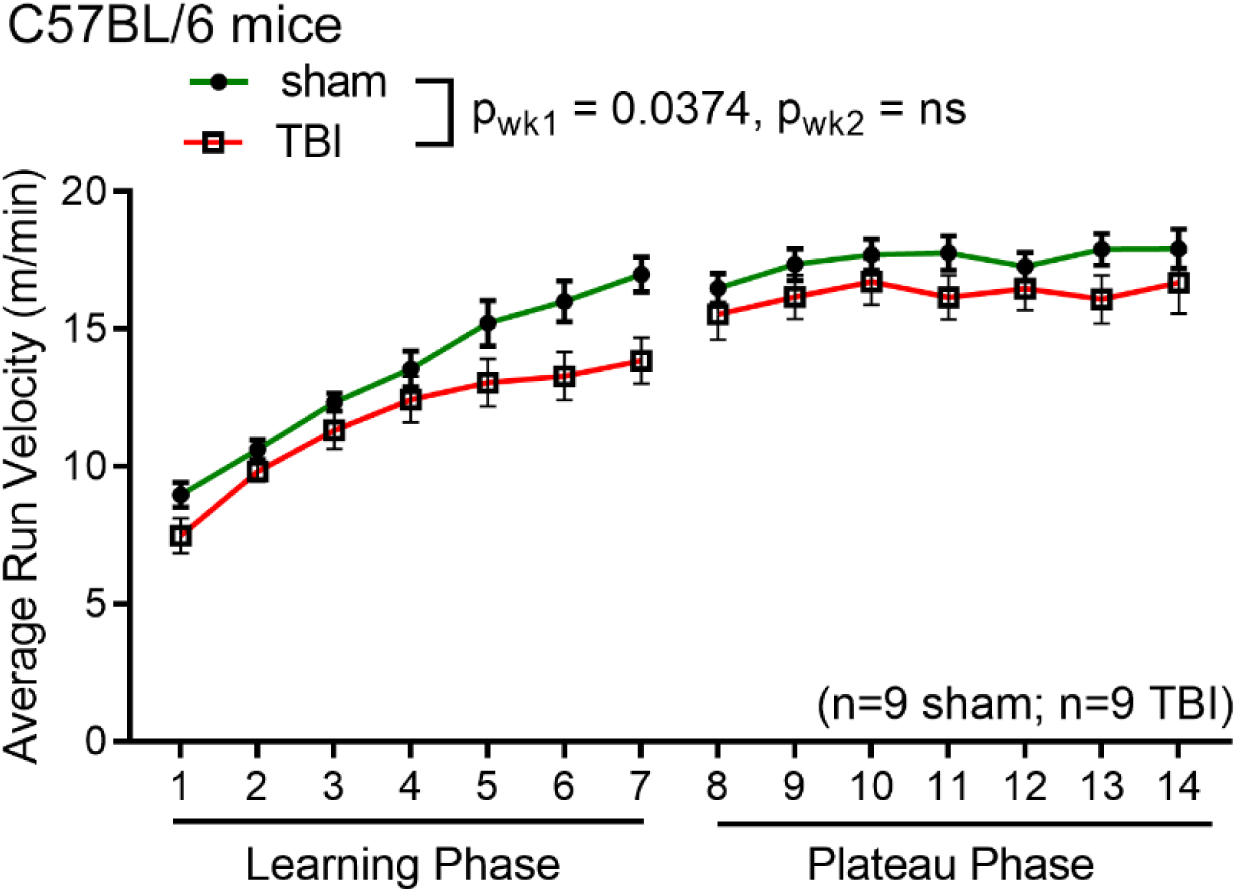
TBI causes a chronic stage motor learning deficit in C57BL/6J mice. Miss-step wheels, which have been shown to engage the CC and be sensitive to myelination status, were used to assess chronic stage changes between sham and TBI mice. The wheels have irregularly spaced rungs to assess motor skill learning (week 1) followed by a plateau phase (week 2) that tests bilateral sensorimotor function. TBI mice show a significant decrease in average running velocity compared to sham during the learning phase (week 1). There was no statistically significant difference between TBI and sham groups during the plateau phase of the assay (week 2). Wheels n = 9 per condition; male C57BL/6J mice (RRID:IMSR_JAX:000664), Jackson Laboratories, Bar Harbor, ME). Two-way repeated measures mixed effects model ANOVA; Learning Phase: Time p < 0.0001, F (2.640, 40.93) = 84.41; Injury p = 0.0374, F(1,16) = 5.151; Time x Injury p = 0.0299, F (6, 93) = 2.457. Plateau Phase: not significant.

**Supplemental Figure S4.**
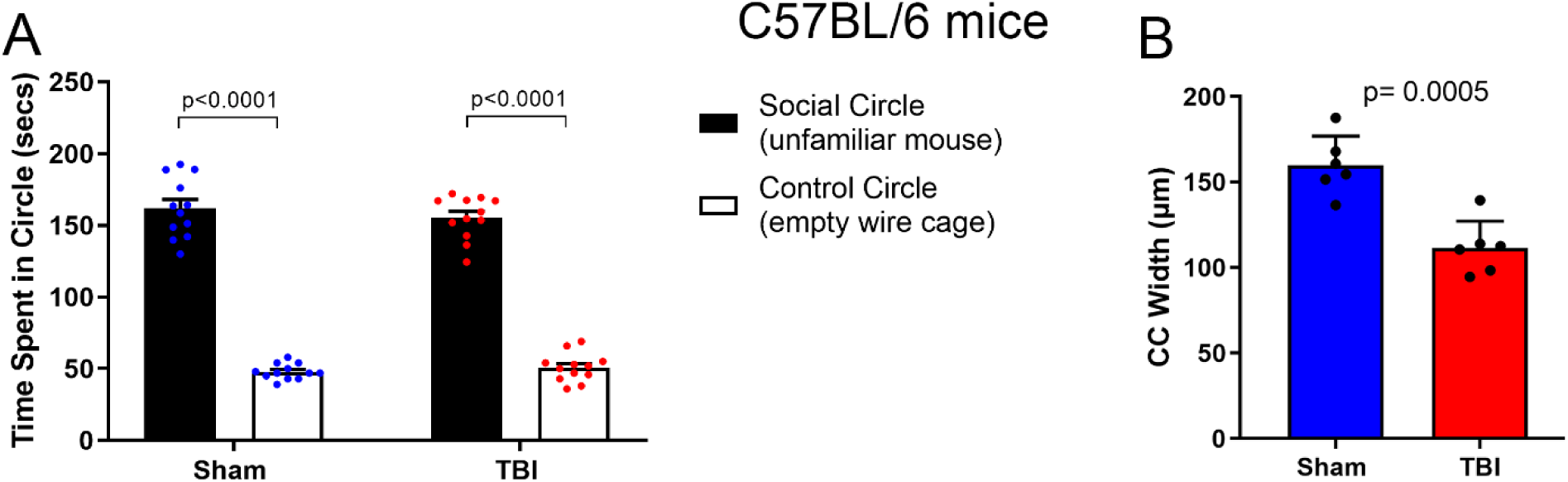
TBI does not induce a late-stage social deficit despite CC atrophy in C57BL/6J mice. In their 8^th^ week post injury mice were tested using a social interaction assay to measure propensity to engage in social behavior. For five minutes mice freely explore a three chamber apparatus with one chamber containing a novel, unfamiliar mouse in a wire carrier and another possessing an empty wire carrier while their time spent interacting with each carrier is measured. Our previous study showed that CC pathology in an experimental model of chronic demyelination induces a social deficit [1]. This study looked to determine if a social deficit, that was not present at more acute time points after single impact TBI [2], develops at a chronic time point. Both TBI and sham mice spent a significantly greater amount of time interacting with the carrier containing the unfamiliar mouse compared to the empty carrier (**A**). There was no statistically significant difference in interaction time between sham and TBI groups. At 10 weeks post TBI mice were sacrificed for neuropathological evaluation of CC width. TBI mice exhibit significant late-stage CC atrophy compared to sham (**B**). The absence of a social deficit at 8 weeks post TBI despite ongoing, chronic atrophy of the CC resulted in the omission of the social interaction assay from our study evaluating the therapeutic benefit of *Sarm1* genetic deletion. Social Interaction n = 12 per condition; male C57BL/6J mice (RRID:IMSR_JAX:000664), Jackson Laboratories, Bar Harbor, ME). Neuropathology n = 6 per condition randomly selected from the MRI cohort. Social interaction: Two-way ANOVA; F(1,44) = 0.2116. Neuropathology: Student’s *t*-test.

